# An Orally Available Cathepsin L Inhibitor Protects Lungs Against SARS-CoV-2-Induced Diffuse Alveolar Damage in African Green Monkeys

**DOI:** 10.1101/2021.07.20.453127

**Authors:** Felix W. Frueh, Daniel C. Maneval, Rudolf P. Bohm, Jason P. Dufour, Robert V. Blair, Kathy Powell, Pyone P. Aye, Nadia A. Golden, Chad J Roy, Sky Spencer, Kasi Russsell-Lodrigue, Kenneth S. Plante, Jessica A. Plante, James McKerrow, Jay Rappaport

## Abstract

The COVID-19 pandemic resulted from global infection by the SARS-CoV-2 coronavirus and rapidly emerged as an urgent health issue requiring effective treatments. To initiate infection, the Spike protein of SARS-CoV-2 requires proteolytic processing mediated by host proteases. Among the host proteases proposed to carry out this activation is the cysteine protease cathepsin L. Inhibiting cathepsin L has been proposed as a therapeutic strategy for treating COVID-19. SLV213 (K777) is an orally administered small molecule protease inhibitor that exhibits *in vitro* activity against a range of viruses, including SARS-CoV-2. To confirm efficacy *in vivo*, K777 was evaluated in an African green monkey (AGM) model of COVID-19. A pilot experiment was designed to test K777 in a prophylactic setting, animals were pre-treated with 100mg/kg K777 (N=4) or vehicle (N=2) before inoculation with SARS-CoV-2. Initial data demonstrated that K777 treatment reduced pulmonary pathology compared to vehicle-treated animals. A second study was designed to test activity in a therapeutic setting, with K777 treatment (33 mg/kg or 100 mg/kg) initiated 8 hours after exposure to the virus. In both experiments, animals received K777 daily via oral gavage for 7 days. Vehicle-treated animals exhibited higher lung weights, pleuritis, and diffuse alveolar damage. In contrast, lung pathology was reduced in K777-treated monkeys, and histopathological analyses confirmed the lack of diffuse alveolar damage. Antiviral effects were further demonstrated by quantitative reductions in viral load of samples collected from upper and lower airways. These preclinical data support the potential for early SLV213 treatment in COVID-19 patients to prevent severe lung pathology and disease progression.

## Introduction

The COVID-19 pandemic has led to a global health crisis of a magnitude not seen since the influenza outbreak of 1918. As of July 2021, > 185 million people have been infected worldwide, almost 4 million have died. In the United States, close to 34 million have been infected with > 600,000 deaths^1^. There is an urgent need for the rapid identification of therapeutic options that limit the pathology caused by SARS-CoV-2 for the management of COVID-19. The current public health response includes the roll out of several vaccines. As of this date, 286 million doses have been given and 130 million individuals are fully vaccinated. While these vaccines are efficacious, the need for an orally available antiviral remains. Viral escape from vaccines and antibody therapies is a major concern. Recent studies demonstrate that variant B.1.1.7 (UK, Alpha) is refractory to neutralization by monoclonal antibodies, and B.1.351 (South Africa, Beta) is highly resistant to neutralization by convalescent and immunized plasma ^2^, and B.1.617.2 (Delta) is highly transmissible and is already accounting for the vast majority of cases world-wide. An oral drug with a host-directed target would be compatible with other drugs in development to create a cocktail to apply selective pressure to minimize new variants, and limit disease spread and severity. Additionally, a host-directed target like cathepsin L acts on highly conserved viral priming mechanisms that are unlikely to undergo significant mutation/modification.

SARS-CoV-2 utilizes binding to a viral Spike (S) protein to enter a host cell. To initiate infection, the S protein requires proteolytic processing (“priming”), mediated by host proteases including cathepsin L, an endosomal cysteine protease. Inhibiting cathepsin L has been shown to prevent SARS-CoV-2 from entering host cells and thereby preventing viral replication by trapping the virus in the endosome for degradation^3^. Blocking coronavirus cell entry through inhibition of a host cell cysteine protease (cathepsin L) function could provide an independent mechanism to limit the pathogenesis and disease severity of SARS-CoV-2. Acute Respiratory Distress Syndrome (ARDS) is the main cause of mortality for COVID-19 patients ^4^. Recent clinical evidence revealed that cathepsin L, and not ACE2, was significantly upregulated in the lungs from COVID-19 patients following autopsy and screening of over 11,000 proteins ^5^. Differential expression of cathepsin L and TMPRSS2 in the upper and lower respiratory tract may account for the difference in efficacy between serine and cysteine inhibitor function^6^ There has also been exploration into a wide range of host protease inhibitors that target other cell-surface serine proteases (e.g., TMPRSS2) and furin^7^. However, many of these inhibitors have off target effects, are difficult to dose in an outpatient setting, or are not compatible with other potential treatments^8–10^. Given the urgent need for orally available antiviral treatments, we are developing SLV213 for the treatment of early-stage COVID-19.

SLV213 (K777) is an orally available cysteine protease inhibitor that acts as a broad antiviral by targeting cathepsin-mediated cell entry. Such entry is required for coronaviruses (e.g., SARS-CoV and SARS-CoV-2, MERS-CoV, NL63) as well as filoviruses (e.g., Ebola-V) for infection. Studies have demonstrated IC_50_ values of K777 consistent with the potential for therapeutic activity inhibiting cathepsins and, thereby, inhibiting infection by coronaviruses ^11^. We previously tested K777 against SARS-CoV-2 *in vitro* using cell lines infected with the coronavirus, demonstrating dose-dependent antiviral effects of K777 in several SARS-CoV-2-permissive mammalian cell lines^3^. Host directed antiviral treatments can be difficult to test *in vitro*, therefore selecting a relevant *in vivo* model is of great importance to demonstrate efficacy of host-directed treatments in a preclinical setting. Typically, non-human primate models have proven to be particularly relevant for COVID-19 because they mimic human disease ^12^. Here we present new data demonstrating prophylactic and therapeutic efficacy of SLV213 in a non-human primate (African green monkey, AGM) COVID-19 model.

## Results

### K777 Pharmacokinetics after Oral Dosing in Non-human Primates

Pharmaceutical development of K777 was previously focused on Chagas’ disease, and extensive preclinical evaluation has been completed to assess the pharmacokinetics, metabolism, and toxicity of K777 in rodents, dogs, and non-human primates^13^. Tolerability and plasma pharmacokinetics of K777 (200 mg/kg) were evaluated in cynomolgus monkeys (n=4) after oral administration. Each animal received 7 daily administrations, and K777 plasma concentration profiles were compared after Day 1 and Day 7. No significant changes in clinical signs, body weight or clinical pathology were observed. Elevations of serum transaminases (AST, ALT, LDH) were measured on Day 3 and Day 8, resolving partially by Day 15 (supplemental data). The maximum plasma concentration (C_max_) was observed 4 hours post-dosing, and there were no apparent differences in pharmacokinetic parameters between Day 1 and Day 7 (Figure 1A). Results in cynomolgus monkeys provided a rationale for the K777 dose and dose regimen to be used in the African green monkey model of COVID-19.

**Figure 1.**
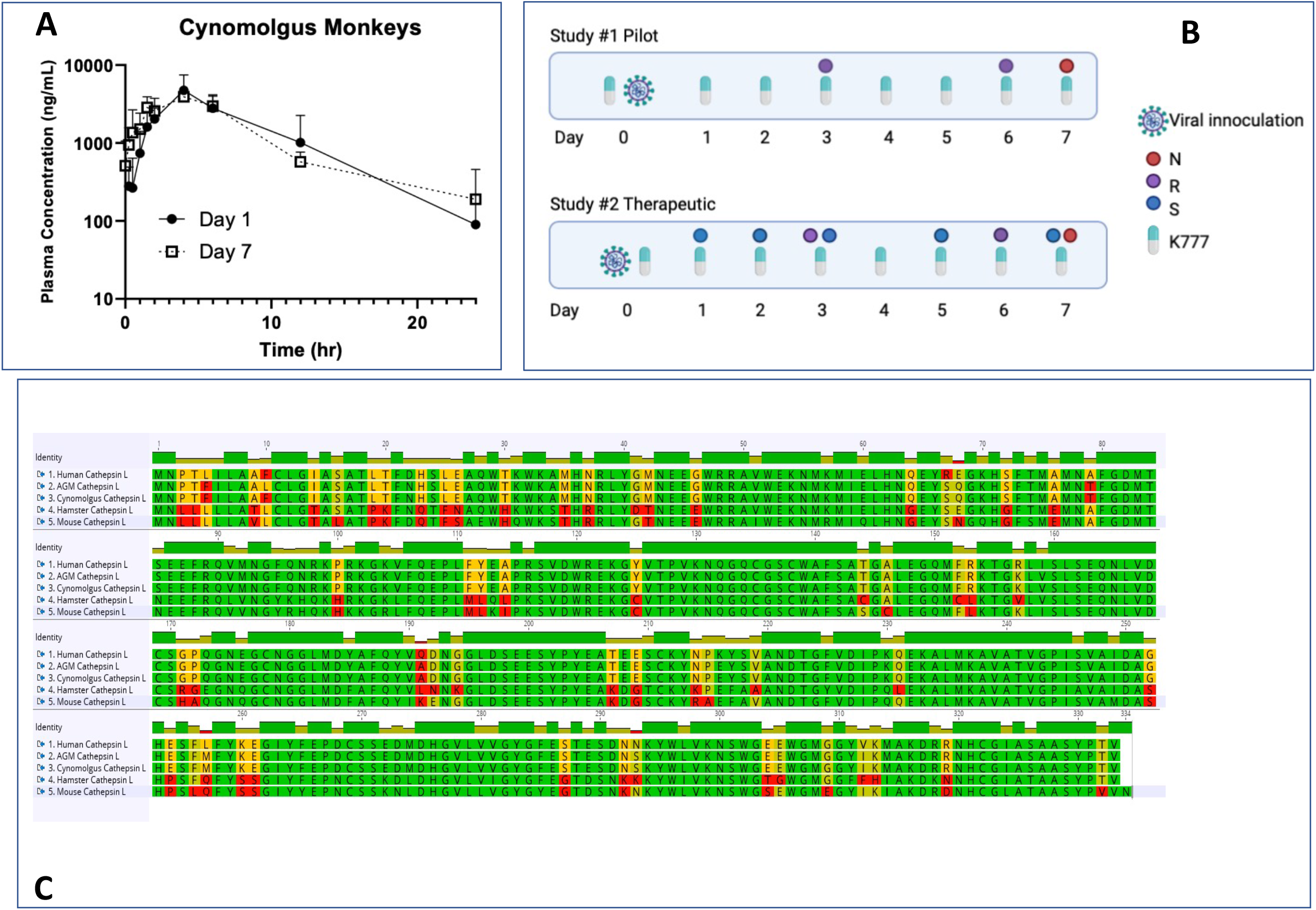
(A) Average (± SD) K777 plasma concentrations in male cynomolgus monkeys after the first (Day 1, closed symbols) and seventh (Day 7, open symbols). (B) Study designs for the Study #1 (pilot study - prophylactic treatment) and Study #2 (therapeutic treatment) in the African green monkey (AGM) COVID-19 model. N denotes necropsy, R denotes chest X-ray, and S denotes samples for viral load (nasal and pharyngeal swabs, BAL fluid). (C) Sequence alignment of AGM, human, cynomolgus, rodent, and hamster cathepsin L.

### African Green Monkey (AGM) COVID-19 Model

Because of similarities to human respiratory anatomy, non-human primates are excellent animal models to investigate the pathogenesis and potential treatments for COVID-19. Serial assessments of blood, tissue, and swab samples after intranasal exposure to SARS-CoV-2 have enabled the translational evaluation of vaccines and therapies for the disease ^14–18^. We have recently reported that the African green monkey (*Chlorocebus aethiops*) model may be particularly effective in modeling ARDS-like clinical effects following exposure to SARS-CoV-2 ^19,20^. Cathepsin L proteases in AGM and human share 96.1% sequence identity, differentiating this animal model from COVID-19 models in other species with less homology in the target protein (Figure 1B).

In this communication, we describe two sequential studies using the AGM COVID-19 model that were designed to evaluate the effects of 7 daily oral doses of K777. Study #1 explored the effects of 100 mg/kg K777 in a prophylactic setting. Results from this pilot study were used to design Study #2, which compared the effects of K777 at two dose levels in a treatment setting. Study designs are shown graphically in Figure 1B.

### Study #1 – Prophylactic Oral Use of K777 Reduced Lung Pathology in AGM Model of COVID-19

Six male young adult African green monkeys (5.75 to 6.7 kg) were enrolled in Study #1 at the Tulane National Primate Research Center (TNPRC). K777 (100 mg/kg, N=4) or vehicle (N=2) was administered by orogastric feeding tube once daily for seven consecutive days. Animals were challenged on Day 1 with live SARS-CoV-2 virus intranasally and intratracheally 2 to 4 hours after K777 dosing. Drug concentrations were confirmed by dosing solutions analysis. Technical issues with daily dosing preparations were noted when they occurred. Quantitative dosing (100 mg/kg) was confirmed on Day 1 and Day 7, but dose preparation errors led to less than expected K777 dosages that varied among animals on Days 2-6. The schedule of assessments and sample collection methods were similar to our prior published work^20^ and is shown schematically in Figure 1B.

Consistent with observations in the cynomolgus monkey, K777 was well-tolerated in African green monkeys. No notable abnormal clinical observations were reported, and there were no significant changes in body weight associated with treatment. Plasma concentration measurements confirmed systemic exposure in K777-treated animals, and average maximum plasma concentrations (C_max_) measured 4 hour post initial dose were 449 ± 197 ng/mL.

Thoracic radiographs (chest x-rays taken on Days 3 and 7 were within normal limits for all animals treated with K777. On Day 3 post infection, both a focal interstitial pattern and a mild bronchial pattern were noted on the radiographs acquired from the two animals (PA51 and PA54) treated with vehicle. These radiographic abnormalities resolved by Day 7 post infection.

Serum chemistry panels demonstrated elevated AST, ALT, GGT, and LDH only in the four K777-treated animals (PA49, PA50, PA52, and PA53). Increased transaminase levels were detected as early as Day 2, consistent with expectations from prior animal studies with K777; however, there was no evidence of liver damage by histopathology at autopsy. Changes in hematology parameters were unremarkable. Gross pathologic findings in all animals were minimal or absent. Pulmonary lesions were present in 5/6 infected animals and bronchial lymphadenopathy (mild to moderate) was noted in all infected animals but appeared more pronounced in untreated monkeys. Gross lesions outside of the respiratory system were only noted in one animal (PA54) and were associated with a prior surgical procedure (peritoneal adhesions).

Lung weight to body weight ratios were significantly lower in K777-treated versus vehicle-treated infected animals (Figure 2A). This measure of efficacy has previously been reported for the antiviral remdesivir in a non-human primate COVID-19 model ^17^. Histopathology was performed on 27 tissues/organs from each of the animals enrolled on this study. Five out of 6 animals exhibited both interstitial pulmonary inflammation and type II pneumocyte hyperplasia, consistent with SARS-CoV-2 associated pneumonia. Representative micrographs are shown in Figure 2B, suggesting a K777-treatment effect.

**Figure 2.**
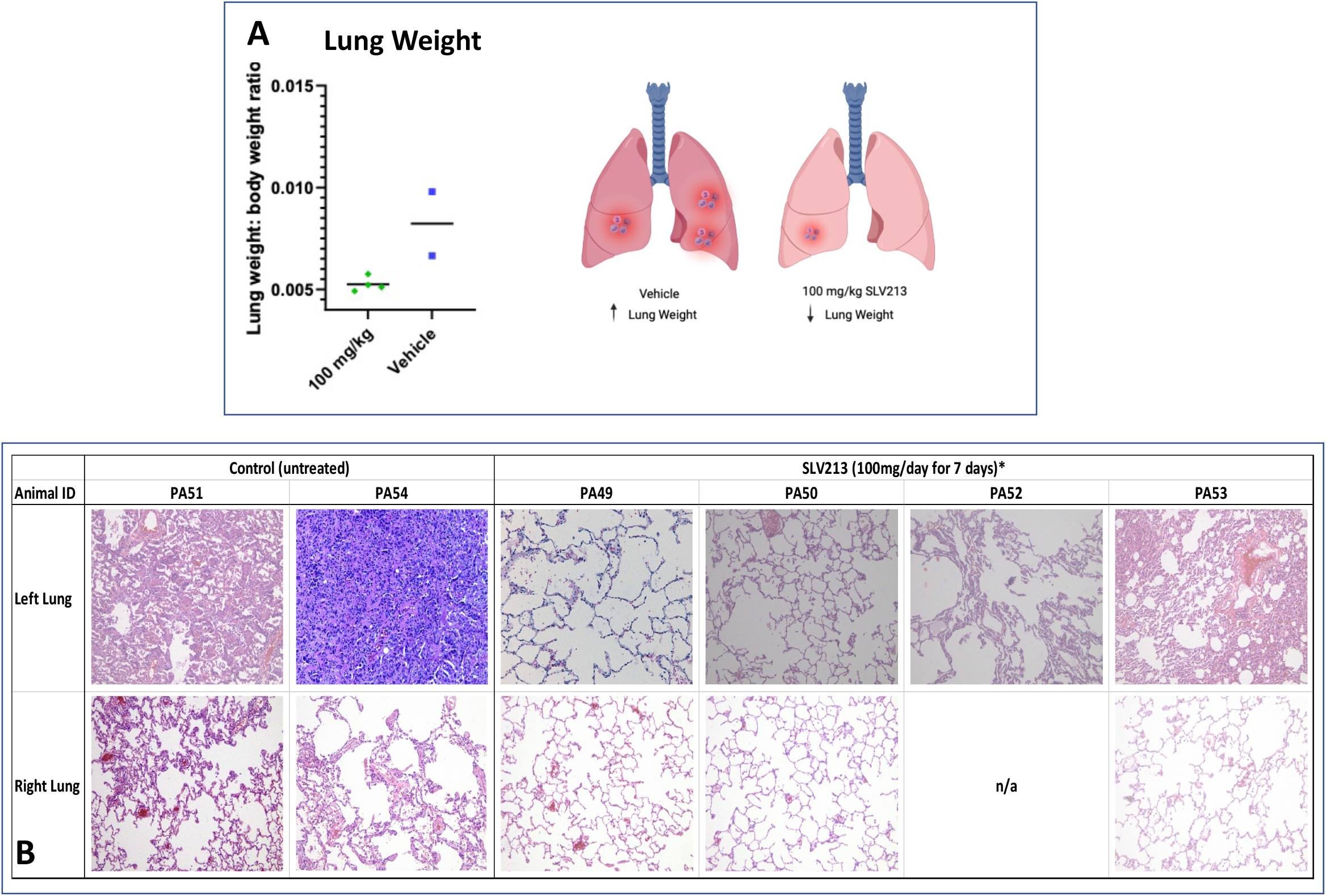
(A) Lung weight to body weight ratio measured in Study #1. Monkeys received vehicle or 100mg/kg K777 by orogastric gavage and necropsy was performed after 7 daily treatments. Ratios were significantly less in K777-treated animals (p<0.05) (B) Representative H&E sections from right and left lungs of each AGM in the study. A reduction in ARDS-like effects and reduction in severity of lesions was noted in K777-treated animals.

In summary, prophylactic oral dosing of K777 was associated with a reduction in SARS-CoV-2-induced lung pathology in the AGM model of COVID-19. Plasma concentrations of K777 and elevated serum transaminase levels were consistent with expectations from prior non-human primate studies. Findings from this pilot study provided a rationale for a second study to evaluate the therapeutic activity of K777 against SARS-CoV-2.

### Study #2 – Therapeutic Oral Use of K777 Reduced Lung Pathology in AGM Model Exposed to SARS-CoV-2

Study #2 was designed to evaluate the effects of oral K777 dosing after exposure to SARS-CoV-2. Like Study #1, animals were challenged with the coronavirus intranasally and intratracheally, however K777 treatment was initiated 8 hours after virus exposure (Figure 1B). Eight young adult male African green monkeys (5.6 to 7.55 kg) received vehicle (N=2) or K777 at 33 mg/kg (N=3) or 100 mg/kg (N=3) by once daily orogastric instillation for 7 days. Expected K777 dosage was confirmed by dose formulation analysis.

Oral dosing of K777 was again well-tolerated in the AGM model. Clinical observations and measurements (e.g., body temp, body weights, SpO2, hematology) were unremarkable throughout the study. Thoracic radiographs (chest x-rays) taken on Days 3 and 7 were within normal limits for all animals on the study, except one vehicle-treated animal (NB79) with enlarged tracheobronchial lymph nodes on both days. Quantitative serum chemistry panels demonstrated K777-treatment related elevations in transaminases (AST and ALT) that increased with dose. Despite elevations in AST and ALT, there was no evidence of liver damage by histopathology at autopsy and no increase in bilirubin levels. Increases in C-reactive protein were detected in all animals, consistent with exposure to SARS-CoV-2 (Figure 3A). Changes in hematology parameters were unremarkable.

**Figure 3.**
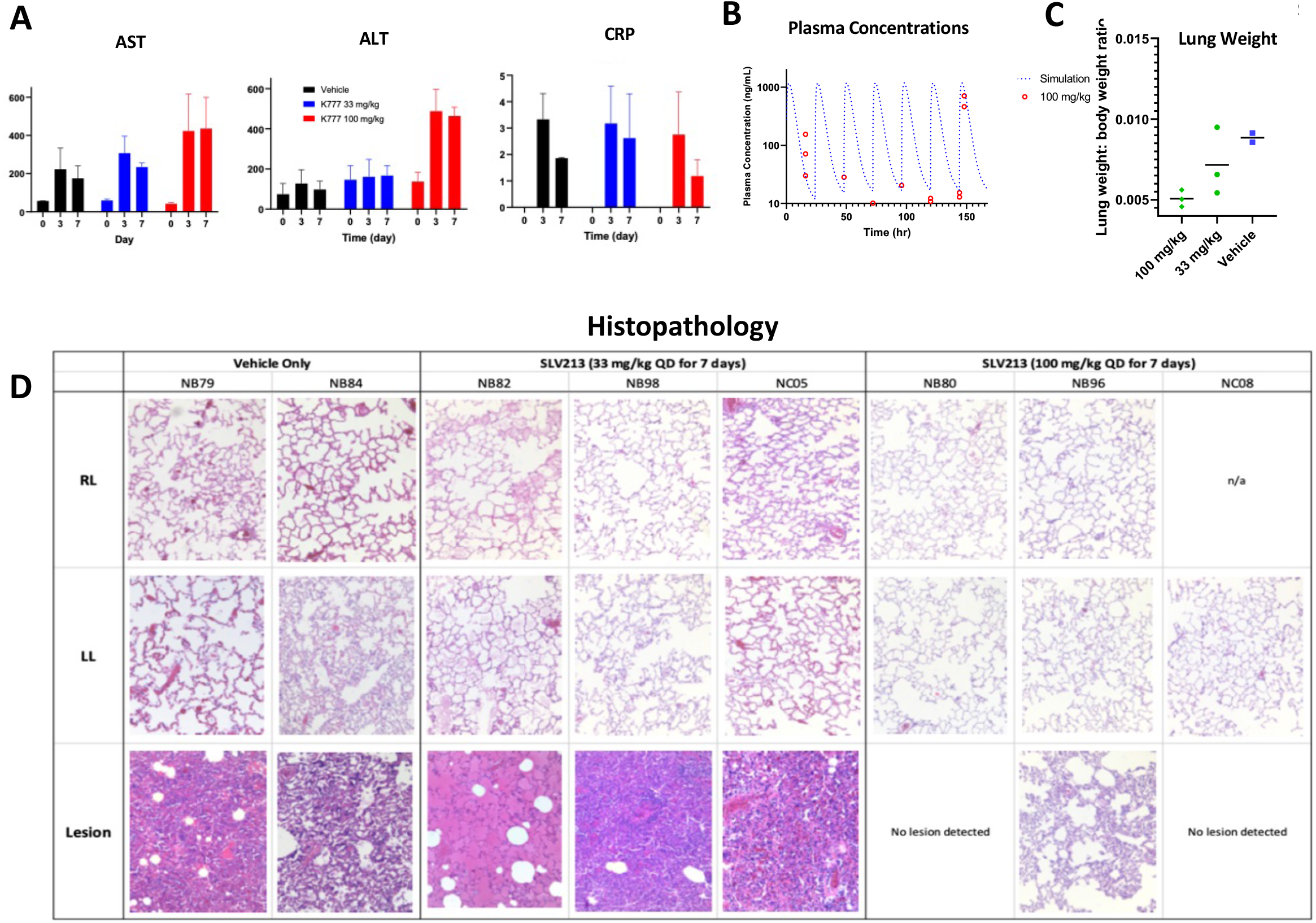
(A) Serum transaminases (AST, ALT) and C-reactive protein (CRP) measured pre-dose, Day 3 and Day 7 in Study #2. Pre-dose CRP values were below the limit of quantification. (B) Simulation of the plasma concentration profile versus the measured plasma concentrations of K777 measured pre-dose (Day 0), 12 hours post initial dose (Day 1), 24-hours post dose (Day 2-5, Day 6 pre) and 4 hours after the last dose (Day 6 post). (C) Lung weight to body weight ratio measured in Study #2. Monkeys received 33 mg/kg or 100 mg/kg K777 or vehicle daily by orogastric gavage. Necropsy was performed after 7 daily treatments. Ratios were significantly less in animals treated with 100 mg/kg vs. vehicle-treated animals (p<0.05) (D) Representative H&E sections from right and left lungs and lesions of each AGM in the study. A reduction in ARDS-like effects and reductions in the severity of lesions was noted in K777-treated animals.

K777 plasma concentrations were measured from blood samples collected prior to daily dosing on Day 0 – Day 6 (C_trough_) as well as 4 hr after dosing on Day 6 (C_max_). C_trough_ measurements demonstrated exposure to K777 after 100 mg/kg dosing without evidence of accumulation after multiple doses, consistent with results reported in cynomolgus monkeys (Figure 1A). C_trough_ values from monkeys treated with 33 mg/kg and all samples analyzed from vehicle-treated animals were below the limit of quantification (10 ng/mL). C_max_ values were dose-dependent and averaged 35.7 ng/mL and 1081 ng/mL after 33 mg/kg and 100 mg/kg dosing, respectively (Figure 3B).

Gross pathologic findings in the lung were noted in 7/8 animals. Gross pulmonary lesions were less severe in K777 treated animals and ranged from absent to mild compared to vehicle controls which were mild or moderate. Pneumonia in K777 treated animals was absent in 1/6 (NB82), minimal in 2/6 (NC08 and NB80), and mild in 3/6 (NB98, NB96, and NC05). Vehicle control animals, in addition to having mild (NB79) and moderate (NB84) pneumonia, both exhibited pleuritis, which was not observed in any of the K777 treated animals. Lung weight to body weight ratios were again significantly lower in K777-treated animals, with a trend toward a dose-dependent K777 effect (Figure 3C), supporting the hypothesis that K777 reduced pulmonary pathology in this model of COVID-19.

Histopathology was performed on 23 tissues/organs from each of the animals enrolled on this study. Pulmonary inflammation and type II pneumocyte hyperplasia was noted within all eight animals and ranged from mild to severe, consistent with our prior experience after SARS-CoV-2 infection. One K777-treated animal (NB80) exhibited mild inflammation and one vehicle-treated animal (NB79) exhibited severe inflammation. All other animals exhibited moderate pulmonary inflammation. Bronchial lymph node hyperplasia was present only in the two vehicle-treated animals (NB84 and NB79) and one K777-treated animal (NC08) all of which had severe type II pneumocyte hyperplasia. Representative micrographs are shown in Figure 3D.

Nasal and pharyngeal swabs were analyzed for SARS-CoV-2 viral genomes to quantify viral load in the upper airway over time. Similarly, bronchoalveolar lavage samples were collected to evaluate coronavirus in the lower airway and lung parenchyma. PCR probes designed to detect different regions of SARS-CoV-2 (sgmE, sgmN, genomic N1) generated similar results. SARS-CoV-2 levels decreased over the 7-day experiment, and K777 tended to reduce these measures of SARS-CoV-2 viral load in both the upper and lower airways (Figure 4).

**Figure 4.**
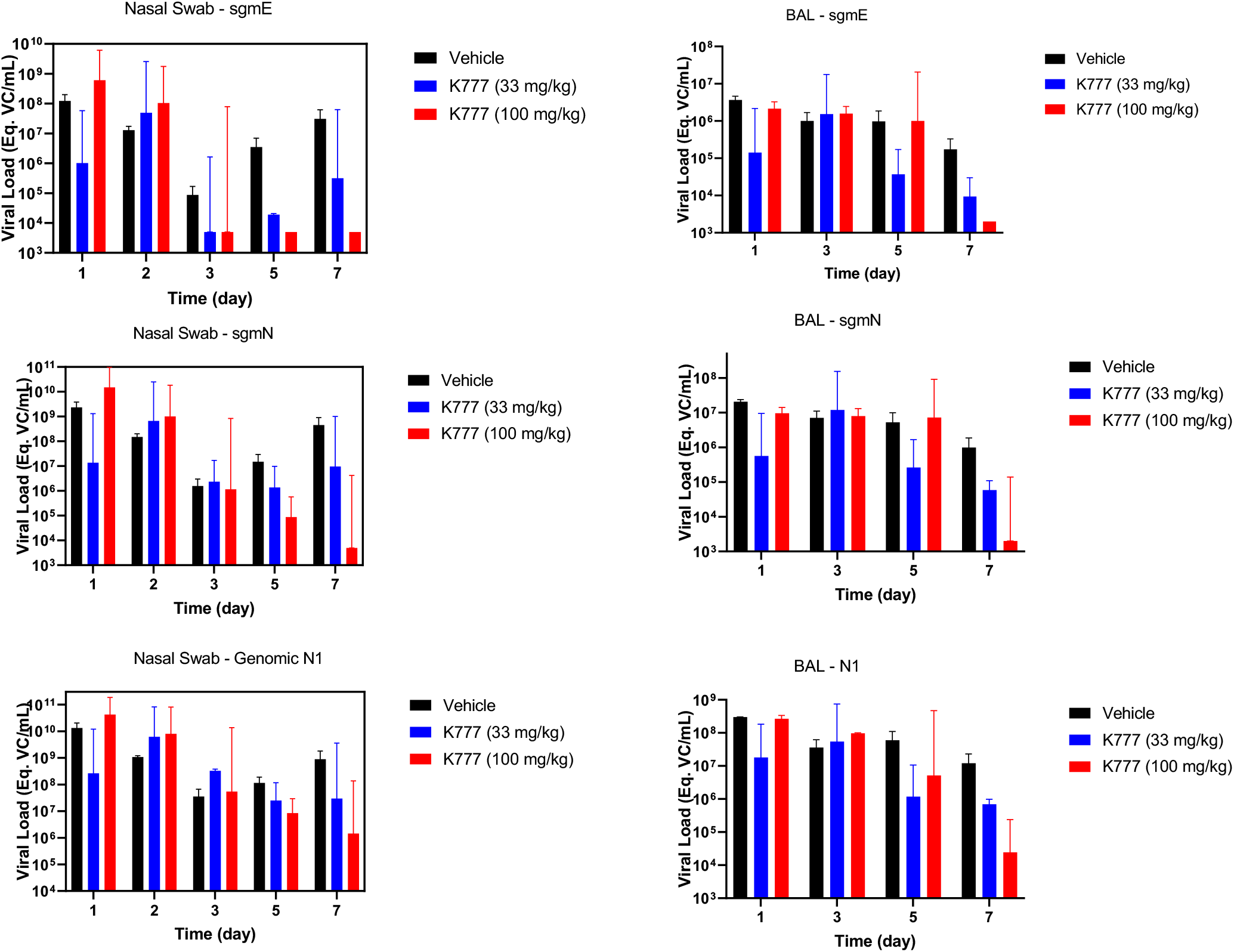
Viral load measurements from nasal swabs (left panel) and BAL fluid samples (right panel) collected in Study #2. PCR-based methods used sequences specific for E and N genes as described in Methods. K777-treatment resulted in a significant reduction in viral load vs. vehicle-treatment on Day 7 (p<0.05).

## Discussion

SLV213, a vinyl sulfone cysteine protease inhibitor, acts as a broad-spectrum antiviral by targeting cathepsin-mediated cell entry. Such entry is required for infection by several different types of enveloped viruses, including coronaviruses, and provides a mechanism-based rationale for investigating SLV213 as a potential treatment against COVID-19. A previous study identified cathepsin L as the target for preventing viral infection of cells ^7^. At the beginning of the pandemic, inhibitors of other host proteins known to be critical for SARS-CoV-2 infection (e.g., ACE2, TMPRSS2) were available for clinical trials ^21,22^. Despite a strong rationale for targeting cathepsin L in the disease ^23^, the absence of an FDA-authorized inhibitor prohibited clinical investigation of this potential therapeutic target in COVID-19 patients. SLV213 is an orally available cathepsin L inhibitor that has recently become available for clinical use and may be ideal for treatment of patients infected with SARS-CoV-2 in an outpatient setting.

An orally available formulation is of utmost importance for treating COVID-19. Current vaccine and antibody treatments are still subject to viral escape mechanisms in less essential regions of the viral genome. A host directed molecule targeting a conserved viral mechanism (i.e., priming, or activation, of the viral Spike protein) limits the likelihood for a viral mutation to escape selective pressure. The only currently available small molecule drug treatment for COVID-19, remdesivir, has a clinical benefit when administered early in disease progression; however, the drug must be administered intravenously due to poor oral bioavailability ^17^. Other drugs such as hydroxychloroquine were promising in *in vitro* models ^24^, but when tested in non-human primates and in patients it did not prove to have added benefit in the treatment of COVID-19 ^25^. Monoclonal antibodies (mAbs) have been approved for treatment in the clinic and are used in tandem with convalescent plasma to reduce disease severity. However, mAbs require parenteral delivery and infusion technology, therefore less than ideal for broad use, particularly in an outpatient setting. Bamlanivimab and Etesevimab were approved in November of 2020 but were short-lived as monotherapies because novel viral variants were quickly escaping the therapy. As of 2021, the Brazil variant P.1 escapes 10-46% of immunity gained from previous infection ^26,27^. Consequently, an orally available treatment that limits the progression of disease and less susceptible to losses in efficacy due to the development of variants is still urgently needed to prevent hospitalizations and avoid overload on medical care facilities, as well as to reduce acute respiratory distress syndrome-related deaths.

As has been demonstrated with other drugs like hydroxychloroquine, *in vitro* assessment of efficacy is not always reflective of clinical effectiveness. We have previously used disease progression in the African green monkey as a translatable model to evaluate other treatments against SARS-CoV-2 ^20^. Non-human primates proved to be useful models to predict their clinical response. African green monkeys were used to model COVID-19 as respiratory infections can sometimes be difficult to model in rodents, (mice and rats do not have a cough reflex) ^28^. Syrian hamsters efficiently replicate virus and display lung lesions ^29^, but compatibility of cathepsin L sequencing homology in that species minimized enthusiasm for use of this model to test K777 (Figure 1C). The design of two sequential studies enabled an efficient evaluation of the potential for K777 safety and efficacy in this non-human primate model.

In this report, we describe dose-dependent effects of K777, an orally available treatment in a non-human primate model of COVID-19. Study #1, despite technical underdosing issues, demonstrated significant positive effects of K777. Because the high K777 dose used in the prophylactic setting of Study #1 showed the potential of K777 as an oral treatment, Study #2 was designed to use K777 in a treatment setting and tested at two different doses. Oral administration of K777 reduced viral load and diminished lung damage caused by SARS-CoV-2, as measured by both gross pathology (lung weight to bodyweight ratio). Histopathology identified diffuse alveolar damage in vehicle but not K777 treated animals. The dramatic reductions in lung weight, indicative of a reduction in consequences of SARS-CoV-2 infection, coupled with the reductions in viral load, suggest cathepsin L is an important target for drug development.

K777 was well tolerated at 33mg/kg and 100mg/kg, with modest elevations in serum transaminases and plasma concentrations that were expected from prior preclinical studies ^13^. These transaminase elevations were transient (returning to normal plasma levels following therapy) and have not been associated with elevations in bilirubin or abnormal liver histopathology^13^. Taken together, these results clearly demonstrate that K777 is safe in preclinical studies and has the potential for an effective oral treatment, in an outpatient setting, for COVID-19 patients.

SLV213 is an orally available treatment likely to reduce the respiratory symptoms associated with clinical progression in COVID-19 patients. African green monkey studies with this drug demonstrated a reduction in lung weight and a reduction in coronavirus-mediated pathology in both prophylactic and treatment settings. The ability to dose individuals early in the disease progression may reduce the number of individuals that progress to severe disease and death as well as reduce the burden on the medical system for intensive care patients. As novel variants emerge the need for therapeutics for the treatment for COVID-19 will increase. Previous infection and immunization are still being evaluated for efficacy as the pandemic is still recent. A Phase I trial with SLV213 has recently been completed and a Phase II trial is planned (https://clinicaltrials.gov/ct2/show/NCT04843787). Expansion of therapeutic agents will only improve outcomes in years to come.

## Methods

### Ethics and Biosafety

The Tulane National Primate Research Center is fully accredited by AAALAC. All animals were cared for in accordance with the NIH’s Guide for the Care and Use of Laboratory Animals. The Tulane University Institutional Biosafety Committee approved the procedures for sample handling, inactivation, and removal from BSL3 containment. The Tulane University Institutional Animal Care and Use Committee (IACUC) reviewed and approved all procedures performed in live animals associated with this study. The study conducted in cynomolgus monkeys was performed at Sierra Biomedical, a division of Charles River Laboratories, Inc. (Sparks, Nevada) according to an IACUC reviewed protocol and standard operating procedures of the laboratory.

### Preparation and Characteristics of K777

For the studies conducted at TNPRC, K777 ((4-methylpiperazine-1-carboxylic acid[1-(3benzenesulfonyl-1-phenethyl-allylcarbamoyl)-2phenylethyl]amide) hydrochloride) was supplied by Selva Therapeutics as dry powder stored at room temperature under desiccant. The test article was formulated in sterile water (10 mg/mL) daily for oral administration. Aliquots of daily dosing solutions were frozen and subsequently analyzed using a qualified HPLC method at KCAS Bioanalytical Services (Kansas City, KS) to confirm dosage. Similar methods were used for the study conducted at Sierra Biomedical to generate a final solution of 20 mg/mL.

### Virus Source and Preparation

The SARS-CoV-2 coronavirus used for the experimental infection at TNPRC was SARS-CoV-2; 2019-nCoV/USA-WA1/2020 (https://www.ncbi.nlm.nih.gov/nuccore; accession number MN985325.1). Virus stock was prepared in Vero E6 cells and sequence confirmed by deep sequencing. Plaque assays were performed in Vero E6 cells. Vero E6 cells were acquired from ATCC (Manassas, VA) [Harcourt].

### Quantification of SARS-CoV-2 Viral Load

Swab samples were collected in DNA/RNA Shield (catalog number R1200; Zymo Research, Irvine, CA) and extracted for viral RNA using the Quick-RNA viral kit (catalog number R1034/5; ZymoResearch). The Viral RNA Buffer was dispensed directly to the swab in the DNA/RNA Shield. A modification to the manufacturers’ protocol was made to insert the swab directly into the spin column to centrifugate, allowing all the solution to cross the spin column membrane. The viral RNA was the eluted (45 mL) from which 5 mL was added in a 0.1-mL fast 96-well optical microtiter plate format (catalog number 4346906; Thermo Fisher Scientific, Waltham, MA) for a 20-mL real-time quantitative RT-PCR (RT-qPCR) reaction. The RT-qPCR reaction used TaqPath 1-Step Multiplex Master Mix (catalog number A28527; Thermo Fisher Scientific) along with the 2019-nCoV RUO Kit (catalog number 10006713; IDTDNA, Coralville, IA), a premix of forward and reverse primers and a FAM-labeled probe targeting the N1 amplicon of the N gene of SARS2-nCoV19 (https://www.ncbi.nlm.nih.gov/nuccore; accession number MN908947). The reaction master mix was added using an X-stream repeating pipette (Eppendorf, Hauppauge, NY) to the microtiter plates, which were covered with optical film (catalog number 4311971; Thermo Fisher Scientific), vortexed, and pulse centrifuged. The RT-qPCR reaction was subjected to RT-qPCR at a program of uracil-DNA glycosylase incubation at 25C for 2 minutes, room temperature incubation at 50C for 15 minutes, and an enzyme activation at 95C for 2 minutes followed by 40 cycles of a denaturing step at 95C for 3 seconds and annealing at 60C for 30 seconds. Fluorescence signals were detected with a QuantStudio 6 Sequence Detector (Applied Biosystems, Foster City, CA). Data were captured and analyzed with Sequence Detector Software version 1.3 (Applied Biosystems).

Viral load was calculated by plotting Cq values obtained from unknown (ie, test) samples against a standard curve that represented known viral copy numbers. The limit of detection of the viral RNA assay was 10 copies per reaction volume. A 2019-nCoV-positive control (catalog number 10006625; IDTDNA) were analyzed in parallel with every set of test samples to verify that the RT-qPCR master mix and reagents were prepared correctly to produce amplification of the target nucleic acid. A nontemplate control was included in the qPCR to ensure that there was no cross-contamination between reactions.

### K777 Pharmacokinetics in cynomolgus monkey

Two male and two female cynomolgus monkeys (2.3 - 3.9 kg) from the Sierra Biomedical colony received 200 mg/kg K777 daily for 7 days by nasogastric gavage (10 mL/kg) in a pilot study to evaluate the toxicity and pharmacokinetics in non-human primates (SRI Study No. M158-01). In-life observations included twice daily cageside observations and daily body weight measurements. Blood samples (approximately 2 mL) for evaluation of serum chemistry and hematology were collected from a femoral vein of all animals pre-study, and on Days 3, 8, and 15.

Whole blood samples (approximately 0.7 mL) were drawn into EDTA tubes for plasma at 0.25, 0.5, 1, 1.5, 2, 4, 6, 12, 24, and 48 hours post-dose on Days 1 and 7. Total K777 plasma concentrations were determined using a qualified method that included liquid/liquid extraction and HPLC analysis with UV detection. Noncompartmental modeling was used to obtain estimates of the pharmacokinetic parameters.

### K777 Treatment in Experimental AGM Model

Wild-caught African green monkeys (*Chlorocebus aethiops*) were quarantined at the TNPRC and were screened for SIV, STLV 1, SRV, and B virus (*Macacine alphaherpesvirus 1*) prior to transfer to a Biosafety Level 3 facility. The quarantine period was shortened for Study #1 (pilot study) and was 90 days for Study #2. Procedures for virus inoculation, daily care of animals, blood draws, nasal swabs and bronchioalveolar lavage (BAL) collection, and chest x-radiography have been previously described (Blair 2021).

Physical exams, body weights, body temperatures, and SpO2 were measured every time animals were accessed. Routine hematology and clinical chemistry were performed pre-dosing and on Day 3 and Day 8 (necropsy), include monitoring of complete blood counts, serum chemistry. Nasal and pharyngeal swabs were collected for viral load on days 1, 2, 3, 5 and 7. BAL samples were collected on days 1, 3 and 5 after placing the animal in a specifically designed chair consisting of a V-trough on a slight recline. A feeding tube was passed into the trachea under direct visualization using a laryngoscope and to a distal subsegmental bronchus. Twenty (20) milliliters of saline solution were instilled into the lung then aspirated and repeated for a second flush. Thoracic radiographs (PA and lateral) were performed pre-study, D3 and D7 using standard techniques at TNPRC.

Orogastric intubation (10 mL/kg) was used to administer the K777 to the experimental animals. 500 mg of K777 was dissolved in 50 mL of sterile water to prepare 10 mg/mL daily dosing formulations. Following administration, orogastric tubes were flushed with 3 mL of physiological solution to ensure the administration of the total volume.

Mucosal fluid secretions were collected throughout the study with swabs which were gently inserted into the nares or pharyngeal regions. Once inserted, the sponge/swab was rotated several times and immediately withdrawn.

Blood samples were collected for K777 plasma concentrations pre-dose on Day 0-6, and 4 hr post-dose on Day 0 (Study 1 only) and Day 6 (Study 2 only). Frozen EDTA plasma samples were analyzed using qualified HPLC-based assays (range 10 - 2000 ng/mL) at KCAS Bioanalytical Services (Kansas City, KS).

Necropsies were performed on all animals on Day 8, seven days post SARS-CoV-2 challenge. Euthanasia occurred following terminal blood collection in deeply anesthetized animals via intracardiac inoculation with sodium pentobarbital. The necropsy was performed by a trained necropsy technician under the oversight of a board-certified veterinary pathologist. Detailed dissection and sampling of the lower respiratory system was performed by a veterinary pathologist as follows: The pluck was removed in its entirety for sampling. The left and right lungs were imaged and then weighed individually. A bronchoalveolar lavage (BAL) was performed on the left caudal lung lobe by instilling and aspirating PBS with a 50 mL syringe. One section from each of the major left and right lung lobes (caudal, middle, and cranial) were sampled fresh. In addition to routine sampling any grossly evident lung lesions were also sampled. Upon completion of sampling the remaining lung tissue was infused with fixative using a 50 mL syringe, and all remaining lung tissue was saved in fixative.

### Statistical Analyses

Graphical analyses were performed with GraphPad Prism software, version 9 (GraphPad Software, San Diego, CA). Test of significance used a standard *t*-test to compare lung weight to body weight ratios.

## Supplemental Information

### Study 1

| **ID** | **Age** | **Body Weight (kg)** | **Treatment** | **Source** |
| --- | --- | --- | --- | --- |
| PA54 | 4 yrs 2 months | 5.75 | Vehicle | Primgen |
| PA53 | 4 yrs 2 months | 6.05 | K777 (100 mg/kg) | Primgen |
| PA52 | 4 yrs 2 months | 5.8 | K777 (100 mg/kg) | Primgen |
| PA51 | 4 yrs 2 months | 6 | Vehicle | Primgen |
| PA50 | 4 yrs 2 months | 6.7 | K777 (100 mg/kg) | Primgen |
| PA49 | 4 yrs 2 months | 6.55 | K777 (100 mg/kg) | Primgen |

### Study 2

| **ID** | **Age** | **Body Weight (kg)** | **Treatment** | **Source** |
| --- | --- | --- | --- | --- |
| NB79 | 8 yrs | 5.8 | Vehicle | Bio Qual Inc. |
| NB80 | 8 yrs | 7.2 | K777 (100 mg/kg) | Bio Qual Inc. |
| NB82 | 8 yrs | 53.9 | K777 (33 mg/kg) | Bio Qual Inc. |
| NB84 | 8 yrs | 5.6 | Vehicle | Bio Qual Inc. |
| NB96 | 8 yrs | 6.35 | K777 (100 mg/kg) | Bio Qual Inc. |
| NB98 | 8 yrs | 7.55 | K777 (33 mg/kg) | Bio Qual Inc. |
| NC05 | 8 yrs | 6.7 | K777 (33 mg/kg) | Bio Qual Inc. |
| NC08 | 8 yrs | 6.05 | K777 (100 mg/kg) | Bio Qual Inc. |

